# Experimental myositis: an optimised version of C-protein-induced myositis

**DOI:** 10.1101/2024.05.18.593723

**Authors:** M Giannini, D Rovito, M Oulad-Abdelghani, N Messaddeq, L Debrut, P Kessler, AL Charles, B Geny, D Metzger, G Laverny, A Meyer

## Abstract

**Introduction:** Inflammatory myopathies (IM) are a group of severe autoimmune diseases, sharing some similarities, whose cause is unknown and treatment is empirical. While C-protein-induced myositis (CIM), the most currently used model of IM, has removed some roadblock to understand and improve the treatment of IM, it has only been partially characterised and its generation limited by reproducibility issues. This study aimed at optimising the generation and the characterisation of CIM.

**Methods:** *In silico* analysis was run to identify the top-3 specific and immunogenic regions of C-protein. The cognate polypeptides were synthetised and used to immunise C57BL/6N mice. Grip strength, walking ability, serum creatine-kinase levels and muscle pathology (histological and electron microscopic features) were assessed. Immune cell proportions and interferon signature in muscles were also determined.

**Results:** Among the three C-protein polypeptides with the highest immunogenic score, amino acids 965-991 induced the most severe phenotype (i.e., 37% decrease in strength, 36% increase in hind base width, 45% increase in serum creatine-kinase level, 80% increase in histological inflammatory score) from day (D) 14 to at least D31 after immunisation [experimental myositis (EM)]. Optical and electron microscopy revealed mononuclear cell infiltrate, myofibre necrosis, atrophy, MHC-I expression as well as sarcolemmal, sarcomeric and mitochondrial abnormalities. Proinflammatory T-lymphocytes, macrophages, type-I and II interferon-stimulated transcripts were found within the muscle of EM mice.

**Conclusion:** EM recapitulates the common hallmarks of IM. This costless, high throughput, reproducible and stable model, generated in the most commonly used background for genetically engineered mice, may foster pre-clinical research in IM.

**Key messages:** *What is already known on this topic:* C-protein-induced myositis is currently the most used model of inflammatory myopathies but has been partially characterised and its generation is limited by reproducibility issues.

*What this study adds.:* Immunisation against the polypeptide encompassing C-protein amino acids 965-991 induces a costless, high throughput, reproducible and stable model of myositis (experimental myositis) that recapitulates the common hallmarks of inflammatory myopathies.

*How this study might affect research, practice or policy:* Experimental myositis, generated in the most used background for genetically engineered mice (C57BL/6N), might foster pre-clinical research in IM.

## Introduction

Inflammatory myopathies (IM) are rare autoimmune diseases which include dermatomyositis, immune-mediated necrotising myopathy, inclusion body myositis, anti-synthetase syndrome and scleromyositis (1–3). Despite their differences, these subgroups share common characteristics (4,5) including proximal weakness, increased serum creatine-kinase (CK) levels and histopathological lesions of the skeletal muscle (i.e. mononuclear cell infiltrates and myofibre abnormalities). Although the severity and the extent of the myofibre lesions vary among subgroups, certain lesions (i.e., major histocompatibility complex [MHC]-I expression, atrophy, necrosis, mitochondrial abnormalities, sarcolemmal irregularities and sarcomeric disruptions) are common to all IM subtypes (6–8). Similarly, although their proportions vary among subgroups, T lymphocytes and macrophage infiltrate (6) along with interferon (IFN) I (9–12) and IFN-II (11–14) signature within the muscle are found in all IM subtypes (15). These findings further indicate that common pathophysiological pathways underpin these diseases (5).

IM are associated with a decreased quality of life and increased mortality (16,17). The aetiology of IM is unknown. Current treatments are empirical (18), partially effective and prone to many side effects. Several mouse models of IM have been established to foster a better understanding of their pathophysiology and to test new candidate drugs (19). Although these models recapitulate some aspects of the disease and have been successfully used to explore preclinical drug efficacy, they have only been partially characterised, and their generation is limited by financial and technical issues.

The most widely used model of IM is C-protein-induced myositis (CIM) (20–24), which is based on a single intradermal injection of a recombinant polypeptide encompassing amino acids (aa) 284-580 of the human fast-type myosin-binding-protein C (MYBPC, hereafter termed C-protein). CIM mice present a decrease in running ability, auto-antibodies targeting C-protein as well as T cell and macrophage infiltrates within the muscle. CD8+ T-cells cytotoxicity is responsible for muscle injury in CIM (25,26). Muscle inflammation peaks at day (D) 21 after immunisation and begins to resolve after D28 (21). However, muscle fibre abnormalities, the detailed composition of the inflammatory infiltrate as well as IFN signature in CIM mice have not been reported.

The CIM model is also limited by both reproducibility and cost. Native C-protein is expensive: several mice are required to extract a protein that only accounts for approximately 2% of the entire amount of muscle proteins, also raising ethical issues (27). Moreover, the purity and immunogenicity of the native protein varies greatly within batches. This could affect the reproducibility of CIM (28). The production of recombinant C-proteins enables a higher throughput, although remains more expensive and less affordable. Reproducibility is also affected by several technical aspects, including difficulty in optimising expression and purification, immunogenicity of tags, etc (29). On the other hand, peptide production is stable, cost-less (100 USD per gram), has a high throughput, and is independent of specific laboratory skills and facilities, hence more reliable. Thus, to overcome these barriers, several groups have attempted to generate IM models using several C-protein polypeptides for immunisation, albeit without fully reproducing CIM (19,28).

In light of the above, the present study aimed at optimising the generation and characterization of the CIM model, taking advantage of an *in silico* analysis to identify the top-3 immunogenic and specific regions of C-protein. Mice immunised with the cognate synthesised polypeptides were deeply phenotyped. The 965-991 aa polypeptide yielded the most severe phenotype [experimental myositis (EM) mice]. Cellular and molecular analyses of the muscle further demonstrated similarities with IM. We thus propose EM as an optimised version of CIM.

## Methods

### Mice

C57BL/6N mice were housed in a temperature- and light-controlled animal facility with food and water ad libitum (Safe Diets, D04, France). Breeding and maintenance were performed according to institutional guidelines. All animal experimental protocols were conducted in an accredited animal house in compliance with French and EU regulations on the use of laboratory animals for research, and approved by the IGBMC Ethical Committee and the French Ministry (#19395-2019022209557705). Blood was collected by inferior palpebral vein puncture; animals were sacrificed by cervical dislocation and tissues were collected and immediately processed for biochemical and histological analysis or frozen in liquid nitrogen.

### *In silico* peptide prediction

The top-3 polypeptides of the human fast-type myosin-binding-protein C were selected by online NHLBI-*AbDesigner* (30), according to Immunogenicity Score (Ig score), Uniqueness Score and Conservation Score.

### Peptide synthesis

Predicted polypeptides were chemically synthesised in-house on an ABI 433A peptide synthesizer using Fmoc chemistry and purified by reverse phase high-performance liquid chromatography using a preparative scale column (Phenomenex: Kinetex EVO C18, 100 A, 5 µM, 250 × 21.2 mm).

### Immunisation

Two hundred µg of polypeptide in 100 µl phosphate-buffered saline (PBS) were emulsified with an equal volume of complete Freund’s adjuvant (FA) (F5881-10 ML, Sigma-Aldrich) and subcutaneously injected on bilateral sides of the hind legs and flanks. Mice injected with the same volume of a saline solution with and without FA were used as controls (ctrl FA and ctrl PBS, respectively). In addition, 2 µg of pertussis toxin (P2980, Sigma-Aldrich) dissolved in 100 µl of PBS were intraperitoneally injected.

### Grip test

Forelimb and hindlimb grip strength was determined by grip Strength Meter (Bioseb). The mean value of three consecutive measurements during the same session was registered as previously described (31).

### Footprint test

The hind feet of the mice were coated with black nontoxic paint (32). Animals were then allowed to walk along a 50-cm-long, 10-cm-wide runway (with 10-cm-high walls) into an enclosed box, as described (32). All mice had three training runs. A fresh sheet of white paper was placed on the floor of the runway for each run. Hind-base width was measured as the average distance between left and right hind footprints. These values were determined by measuring the perpendicular distance of a given step to a line connecting its opposite preceding and proceeding steps as previously described (32).

### Histochemistry

Muscle specimens (gastrocnemius-soleus, quadriceps, tibialis anterior) were rapidly frozen in isopentane cooled in liquid nitrogen and maintained at –80°C until use. Serial 10-µm transverse muscle sections were obtained with a cryostat (LEICA CM3050S) for histochemical staining. Sections were stained with haematoxylin-eosin (H&E) and NADH tetrazolium reductase, as described (31). Necrosis was defined by pale and/or hyalinised staining on H&E (33). Regenerating fibres were identified by increased basophilia on H&E-stained sections and/or centronucleated fibres.

### Microscopic acquisition

Images were acquired using an upright motorised microscope (Leica DM 4000 B) fitted with the CoolSNAP HQ2 (Photometrics) and the Micro-Manager software. Fiji software was used for image editing (31).

### Histological inflammation grading

Inflammation in histological muscle sections was graded as described (34,35). Briefly, grade 1 was assigned for involvement of <5 muscle fibres; grade 2 for a lesion involving 5–30 muscle fibres; grade 3 for a lesion involving a muscle fasciculus; and grade 4 for diffuse and extensive lesions. Three sections from each block were assessed and the lesion with the highest grade was selected to calculate the average score. When multiple lesions with the same grade were found in a single sample, 0.5 point was added to the grade.

### Necrotic fibres quantification

Quantification of necrotic fibres was performed as described (36). Briefly, H&E-stained slides were scanned using the NanoZoomer digital slide scanner (Hamamatsu) and analysed using the NDP.view2 Viewing software (Hamamatsu). The number of necrotic fibers in three muscle sections per animal was determined and averaged.

### Cross-sectional area (CSA) and fibre diameter assessment

TRITC-conjugated wheat germ agglutinin was used for membrane staining (W32464, Thermo Fisher) (37) at 1/500 dilution after paraformaldehyde (PFA) fixation (4%), Triton-X100 0.5% permeabilisation and blocking of non-specific binding by incubation in 5% non-fat dry milk solution (38). Slides were scanned in fluorescence mode and analysed as described above. Myofibre size, cross-sectional area (CSA) and fibre diameter were determined. The minimum Feret diameter (minFeret) was assessed since the latter is the least affected by distortion due to oblique cross-sectioning of muscle tissue (39). Muscle fibres were segmented using Cellpose (40), an artificial intelligence-based program. The pre-trained “cyto” model was selected with a diameter set to 50 pixels and the regions of interest (ROI) generated by Cellpose were saved in a Fiji (41) compatible format. Thereafter, the ROI manager of Fiji was used to perform the minimum Feret radius and surface area measurements.

### Electron microscopy

Ultrastructural analyses were performed as previously described (31). Skeletal muscle samples were fixed by immersion in 2.5% glutaraldehyde and 2.5% paraformaldehyde in cacodylate buffer (0.1 M, pH 7.4), washed in cacodylate buffer for 30 min and stored at 4°C. Post-fixation was performed with 1% osmium tetraoxide in 0.1 M cacodylate buffer for 1 h at 4°C and dehydration through graded alcohol (50, 70, 90 and 100%) and propylene oxide for 30 min each. Samples were oriented longitudinally and embedded in Epon 812. Ultrathin sections were prepared at 70 nm and contrasted with uranyl acetate and lead citrate, and examined at 70 kv with a Morgagni 268D electron microscope. Images were captured digitally by a Mega View III camera (Soft Imaging System).

### Blood analysis

Blood collected at sacrifice was maintained overnight at 4°C and centrifuged at 400 g for 10 min at 4 °C. Serum was stored at -80 °C.

Creatine-kinase (CK) activity was determined (Creatine Kinase Activity Assay Kit, (MAPK116-1KT, Sigma-Aldrich) in serum according to the manufacturer’s protocol.

### Muscle RNA extraction and RT-qPCR analysis

Total RNA was isolated from gastrocnemius muscle using TRIzol reagent (Invitrogen) according to the manufacturer’s instructions. RNA was quantified by spectrophotometry (Nanodrop, Thermo Fisher). cDNA was synthesised from 2 μg RNA by reverse transcription using random primers and SuperScript IV reverse transcriptase (Invitrogen, Life Technologies), according to the manufacturer’s protocol. Quantitative RT-PCR analysis was performed with selective primers (Supplementary Table 1) using the Light Cycler 480 SYBR Green I Master X2 Kit (Roche), according to the manufacturer’s protocol. For each sample, the relative abundance of the transcripts of a given gene was normalised to those of a housekeeping gene (18S).

### Flow cytometry

#### Tissue dissociation

Muscle dissociation into single cells was performed following a previously described protocol (42) with minor modifications. Briefly, muscles (gastrocnemius, soleus, tibialis anterior and quadriceps of both legs) were dissected, mechanically minced, and enzymatically dissociated in Ham’s F-10 (Hyclone) supplemented with 10% horse serum (Life Technologies), 100 units/ml penicillin and 100 µg/ml streptomycin (Omega Scientific) containing 1 mg/ml collagenase type I (17018029 Thermo Fisher Scientific/Gibco) and 5 U/ml dispase (#07913 STEMCELL Technologies) with mild agitation for 60 min in a water bath at 37°C. Cellular clumps were dissociated into single cells by passing the suspension at least 10 times in 5 and 10 ml serological pipettes, followed by 10 times through a 20-gauge needle and a 70-μm cell strainer.

#### Immunophenotyping experiments

The repertoire of mononuclear cell subpopulations in muscle was established by flow cytometry. Cell suspensions isolated from muscle were incubated with an antibody mixture (Supplementary table 2) for 40 minutes on ice. Among DAPI-negative cells (living cells), haematopoietic cells were identified by CD45 marker. Among CD45+ cells, lymphocytes were gated according to side scatter, based on their relative size. Two lymphocyte populations were then subsequently gated: CD3+ (T-lymphocytes) and CD3-lymphocytes. Among CD3+ cells, CD4+, CD8+ and double negative cells (CD3+CD4-CD8-) were identified. From CD45+ cells, CD11b+ cells were gated to select macrophages (F4/80+ cells). Among F4/80+ cells, Ly6C+ and CD206+ macrophages were selected. Data of individual mice were acquired using a BD LSRFortessa Cell Analyzer and analysed using FlowJo software.

### Data analysis

No inclusion/exclusion criteria, and no method of randomization were used in this study. No blinding was used for animal studies.

The normality of the distributions was verified with the Shapiro-Wilk test. Continuous variables were summarized as median and interquartile range or mean + standard error of mean (s.e.m.) according to their distribution. Statistical comparisons of data between two groups were made by a Student’s t-test and those between three and more by one-way ANOVA followed by a post-hoc analysis (Tukey’s test). Data were considered to be statistically significant if p< 0.05 and are indicated by * or $ in the figures.

Statistical significance was set at p-value <0.05. Statistical analyses were performed using GraphPad Prism 9. The exact significant p-values are provided in figures caption.

## Results

### Amino acids 663-684, 866-890 and 965-991 are the top three potential immunogenic regions of C-protein in mice

The online NHLBI-AbDesigner software was run to identify specific and immunogenic regions of C-protein, encoded by *MYBPC2* (30). The top 3 polypeptides fulfilling the following criteria were selected: i) the highest antigenicity score (immunogenicity); ii) the lowest homology with other MYPC isoforms or other proteins (uniqueness); iii) the lowest conservation between mouse and human isoforms to increase peptide immunogenicity in mice (conservation); iv) the absence of aa susceptible to post-translational modifications. Peptide 1 (aa 663-684), peptide 2 (aa 866-890) and peptide 3 (aa 965-991) are presented in Figures 1a, b. Peptides 2 and 3 are located within one of the two most immunogenic regions of C-protein (fragment 4) identified by Sugihara et al (21).

**Figure 1.**
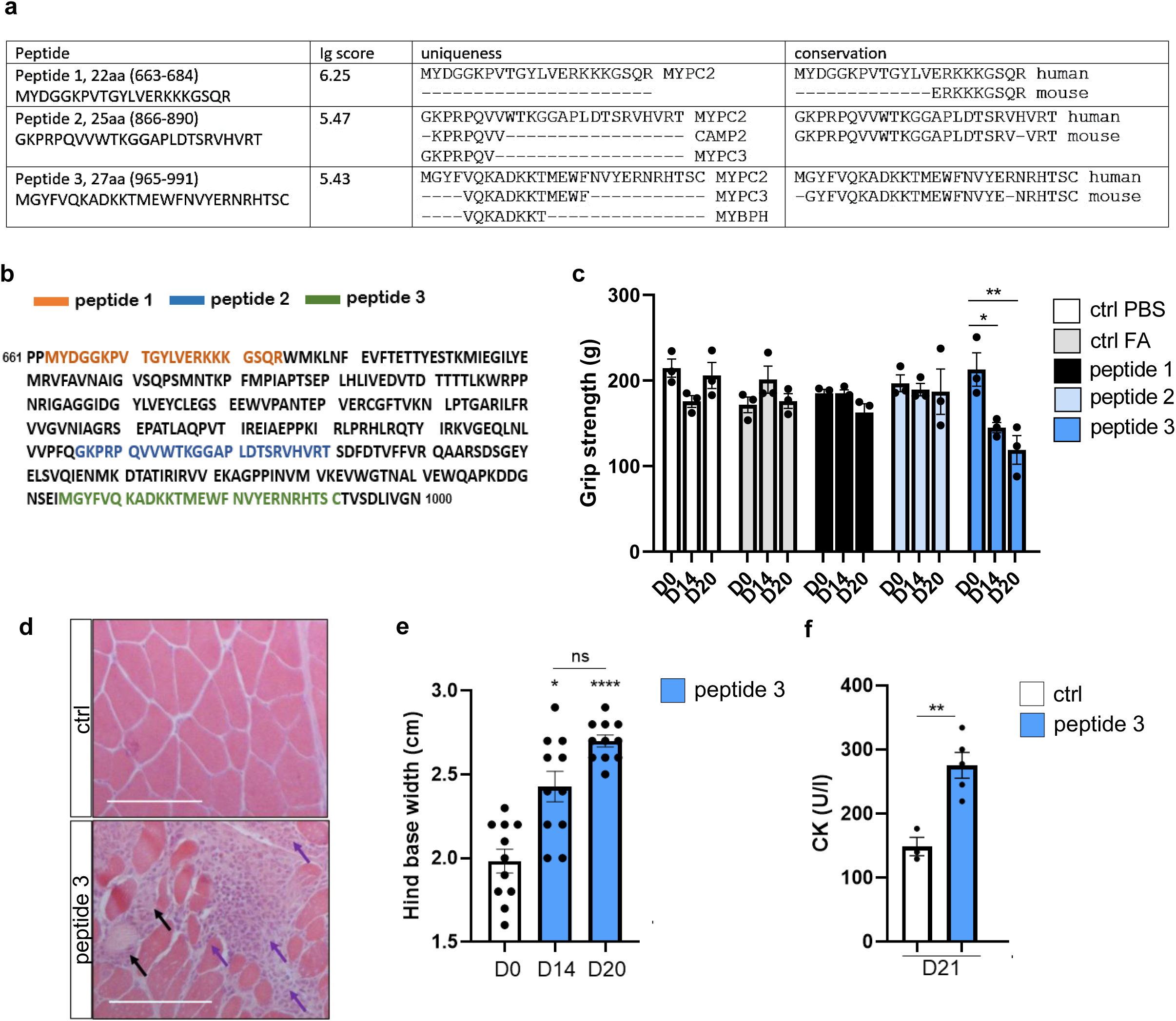
Muscle weakness, walking impairment and muscle lesions in mice treated with polypeptides of human fast-type myosin-binding protein C (MYPC). a) The predicted features of three polypeptides according to NHLBI-AbDesigner are summarised, in particular Immunogenicity Score (Ig score), Uniqueness Score and Conservation Score. Amino acid sequence of human fast type MYPC. b) The three polypeptides used for immunisation are identified in coloured text. c) Grip strength in a cohort of male mice treated with the three C-protein peptides. Mice treated with PBS alone (ctrl PBS) or with an emulsion of PBS and CFA (ctrl FA) were used as controls (ctrl). Muscle strength is expressed as mean ± standard error of the mean. n=3 per group. *: p= 0.04, **: p= 0.001 versus day (D) 0. d) Haematoxylin and eosin (H&E) staining of gastrocnemius muscle sections from peptide 3-treated and control male mice at D21. Necrotic fibres (pale fibres) and immune cell infiltrations are indicated by black and purple arrows, respectively. Scale bar: 100 µm. e) Footprint test performed in female mice treated with peptide 3 at D0, D14 and D20. Hind base width was measured (cm) and expressed as mean ± standard error of the mean (n= 11). *: p= 0.03, ****: p< 0.0001. f) Serum creatine kinase (CK) levels at D21 in female mice immunised with peptide 3 compared to ctrl. **: p= 0.005.

### Immunisation of mice with the synthetic polypeptide encompassing aa 965-991 of C-protein induces muscle weakness, an increase in CK levels and muscle histopathological lesions common to IM

The production and purification procedure yielded polypeptides of molecular weights of 2497.90 mg/mol (peptide 1), 2743.17 mg/mmol (peptide 2) and 3369.86 mg/mmol (peptide 3) with a purity of 70.4% (peptide 1), 92.7% (peptide 2) and 93.4% (peptide 3), respectively (Supplementary Figure 1a).

To compare the ability of these peptides to induce myositis, eight- to ten-week-old mice were immunised with a single subcutaneous injection (on both sides of the hind legs and flanks). Mice administered with complete Freund’s adjuvant (FA) + peptide 3 exhibited a progressive decrease in grip strength from D14 (-32%, p=0.04) to D20 (-44%, p=0.001) after immunisation (Figure 1c). Analysis of H&E stained cryosections from gastrocnemius muscle harvested at D21 revealed mononuclear cell infiltration and necrosis of muscle fibres (Figure 1d). By contrast, FA + peptide 1 and FA + peptide 2 did not affect muscle strength and no histological lesion was found in muscle (Supplementary Figure 2a). Immunization with peptide 3 hence led to the most severe phenotype (experimental myositis, EM). Given that FA with phosphate-buffered saline (PBS) (ctrl FA) had no effect on muscle strength and histology (Figure 1c, Supplementary Figure 2a), ctrl PBS mice were used as controls (ctrl) in the subsequent experiments.

Further characterisation of EM mice highlighted an increase in hind base width at D14 (+22%, p=0.03) and D20 (+36%, p=0.0001) (Figure 1e; Supplementary Figure 2b). Moreover, serum CK activity at D21 was 2-fold higher in EM mice than that in ctrl mice (p=0.005) (Figure 1f). Concurrently with a 23% decrease in muscle strength between D31 and D0 (p=0.0001) (Figure 2a), cryosections of the tibialis anterior, gastrocnemius and quadriceps muscles revealed inflammatory infiltrates, necrotic myofibres and numerous small rounded myofibres at D14, D21 and D31 after immunisation (Figure 2b; Supplementary Figure 3a). A more detailed analysis of the gastrocnemius muscles at D21 revealed that the mononuclear cell infiltrates scored 6-fold higher than in controls (p=0.004) and were located in the endomysial, perimysial and perivascular regions (Figure 2c). The number of necrotic fibres in EM mice was greater than ten times higher than in controls (p=0.01) (Figure 2d). Fibres CSA and minFeret diameter were reduced by 36% (p=0.03) and 23% (p=0.03), respectively, and the percentage of atrophic fibres (CSA less than or equal to1000 µm^2^) was 2.4-fold higher (p=0.005) in EM mice than in ctrl mice (Figure 2e, 2f, 2g, 2h). Sarcolemmal expression of MHC-I was found in EM mice, but not in ctrl mice (Figure 2i). NADH staining revealed mitochondrial abnormalities, such as linearisation of sarcoplasmic membranes and motheaten fibres (Figure 2j). Electron microscopic analysis of gastrocnemius (Figure 2k) and quadriceps (Supplementary Figure 4) muscles revealed additional lesions of the myofibres including i) sarcolemmal invagination, ii) sarcomeric disruption and Z-lines misalignments, iii) T-tubule dilatations, iv) mitochondria fragmentation.

**Figure 2.**
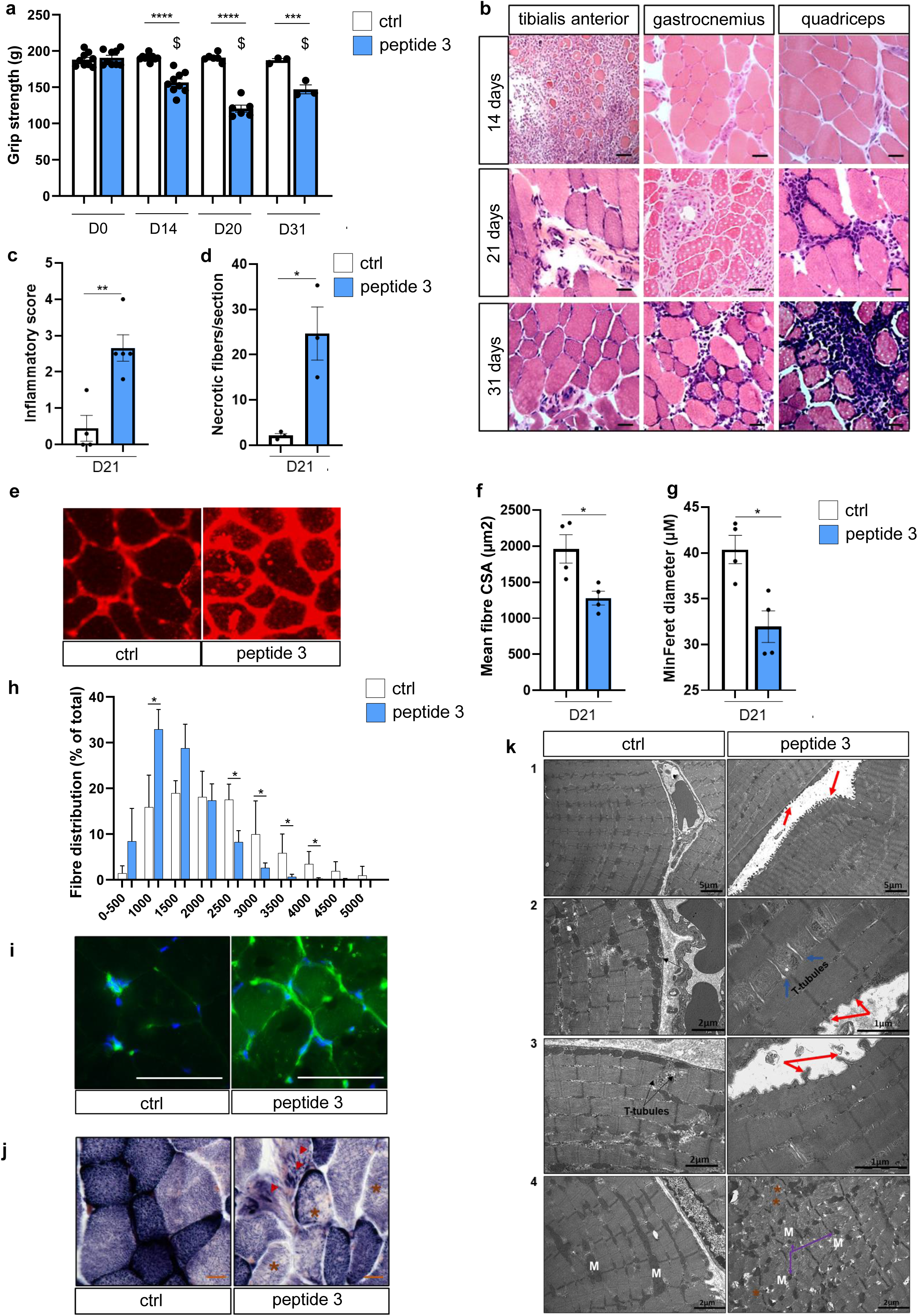
Characterisation of muscle impairment in female mice treated with a peptide encompassing amino acids 965-991 (peptide 3) of fast type skeletal muscle C-protein. a) Grip strength measurements in mice at day 0 (D0), 14 (D14), 20 (D20) and 31 (D31) after subcutaneous (SC) administration of the peptide 3. Mice injected with PBS were used as ctrl. Muscle strength is expressed as mean ± standard error of the mean. n≥ 3 per group. ****: p< 0.0001, ***: p= 0.0003 compared to ctrl at the same timepoint; $: p< 0.0001 versus grip strength at D0 in peptide 3-treated mice. b) H&E staining of tibialis anterior, gastrocnemius and quadriceps cryosections at D14, D21 and D31 after immunization with peptide 3. Scale bar: 50 µm. c) Histological inflammatory infiltrate score in H&E-stained slides from control and peptide 3-treated mice at D21. Each dot represents the average score of three sections for each mouse (n= 4 for each condition). d) Necrotic fibres quantification in H&E-stained sections from control and peptide 3-treated mice at D21. The number of necrotic fibres in three muscle sections per mouse (n= 3 for each condition) was determined and averaged. e) Wheat germ agglutinin TRITC-conjugated staining in gastrocnemius cryosections from control and peptide 3-treated mice at D21. Slides were scanned in fluorescence mode and analysed to determine fibre size. Cross-sectional area (CSA) (f) and minimum Feret diameter (minFeret) (g) measurements on gastrocnemius cryosections from control and peptide 3-treated mice at D21. *: p= 0.03. Results are expressed as mean ± standard error of the mean. Each dot represents the average results for one mouse (n= 4). h) Size distribution of 16040 and 20111 fibres from the gastrocnemius muscle of mice treated with peptide 3 (n= 4) and ctrl mice (n= 4), respectively, at D21. *: p< 0.05. i) Major histocompatibility complex-I staining (green) in gastrocnemius muscle from control and peptide 3-treated mice (n= 3 for each condition) at D21. Nuclei: DAPI (blue). Scale bar: 50 µm. j) Mitochondrial abnormalities assessed by reduced nicotinamide adenine dinucleotide-tetrazolium reductase (NADH) staining in gastrocnemius muscle at D21. Motheaten fibres (brown asterisk) and linearisation of sarcoplasmic membranes (red arrowheads) are shown (n= 3 for each condition). k) Electron microscopy analysis of gastrocnemius muscle at D21 in control and peptide 3-treated mice (n= 3 for each condition). Sarcomeric disruption (panel 4), Z-lines misalignments (brown asterisk), T-tubule dilatations (light blue arrows), sarcolemmal invagination (red arrows) and mitochondrial (white M) fragmentation (violet arrows) are shown in peptide 3-treated mice. Representative images are shown.

Taken together, these results demonstrate that EM mice recapitulate common clinical, biological and pathological hallmarks of IM.

### Muscle from experimental myositis mice displays a pro-inflammatory environment resembling IM

To further assess the similarities between EM and IM at the cellular and molecular levels, a comprehensive analysis of the immune compartment and measurement of interferon-stimulated transcripts were performed in muscle of EM mice at D21.

Flow cytometry analysis (Supplementary Figure 5) revealed that the proportion of CD45+ cells (immune cells) was 12-fold higher in EM mice than in ctrl mice [16.8 (10.5-19.4) vs. 1.4% (0.8-5) of singlets, p= 0.04] (Figure 3a).

**Figure 3.**
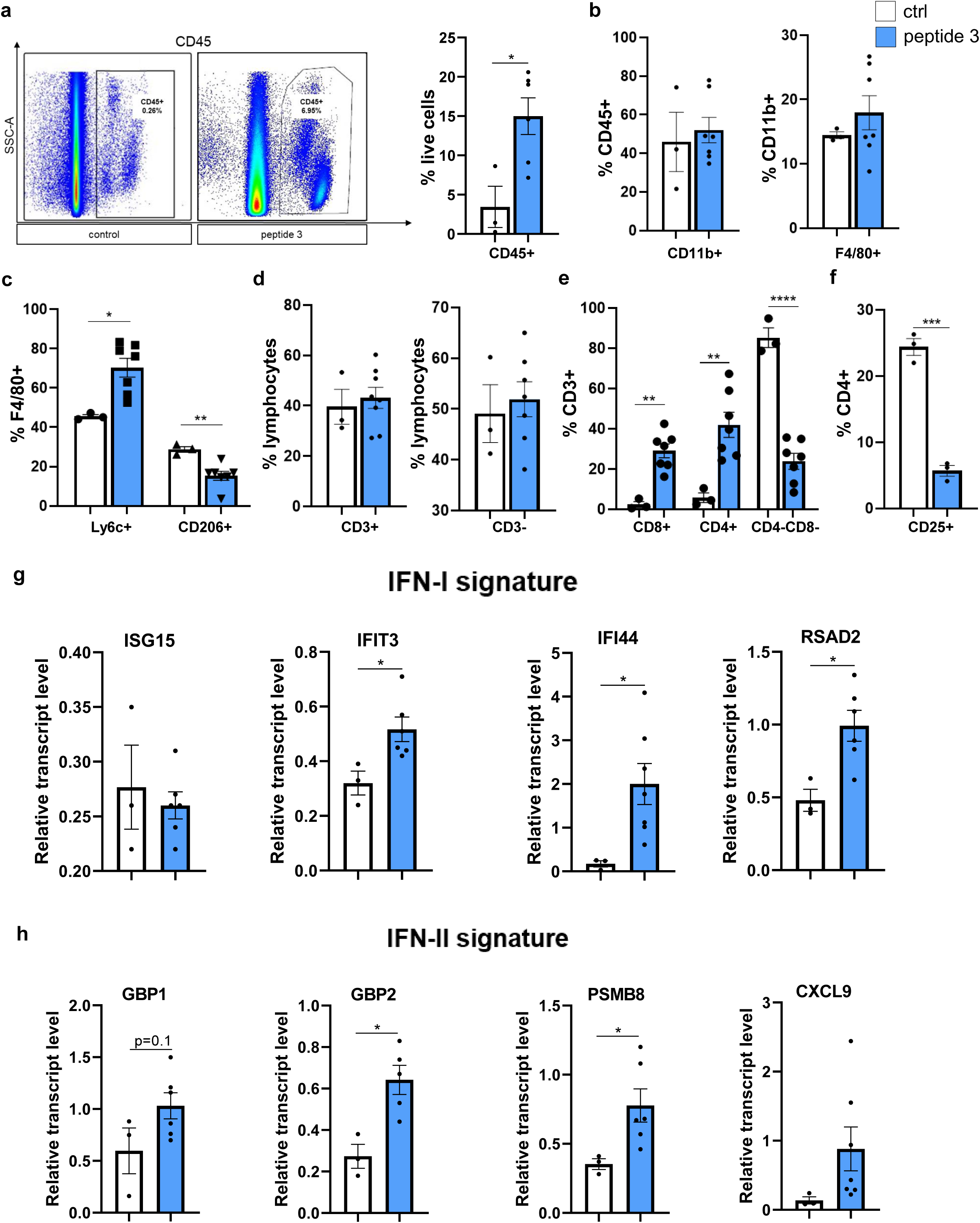
Characterisation of the muscle proinflammatory environment in female mice treated with a peptide encompassing amino acids 965-991 (peptide 3) of human fast-type MYPC. Flow cytometry analysis performed on muscle (anterior tibialis, gastrocnemius, quadriceps) from peptide 3-treated mice (n≥ 3) and control mice (n≥ 3) from independent cohorts: a) representative image of the gating strategy used to select CD45+ cells (immune cells) and lymphocytes. *: p= 0.02. b) Myeloid compartment. c) Macrophage distribution. *: p= 0.01; **: p= 0.007. d) Lymphocytes distribution. e) T-lymphocytes distribution (CD3+ cells). **: p< 0.01; ****: p< 0.0001. f) Regulatory T-cells (CD4+ CD25+ cells). ***: p= 0.0002. Each bar represents the mean ± standard error of the mean. g, h) Relative transcript levels of (g) interferon (IFN) I and (h) IFN-II signature in gastrocnemius muscle of control and peptide 3-treated mice (n≥ 3 independent biological replicates for each condition). The levels were normalized to the 18S gene (housekeeping). *: p< 0.05.

The proportion of CD11b+ cells (myeloid cells) and F4/80+ cells (macrophages) were similar in EM and ctrl mice (Figure 3b). Nevertheless, the proportion of F4/80+Ly6c+ cells (pro-inflammatory macrophages) in EM mice was 1.7-fold higher [76.1 (56.6-81.8) vs. 45.1% (44.1-47.3), p=0.01] while that of F4/80+CD206+ cells (anti-inflammatory macrophages) was 1.7-fold lower [16.8 (14.5-16.9) vs. 29% (26.2-31.2), p=0.007] (Figure 3c). The proportion of CD3+ (T lymphocytes) and CD3-cells was similar in EM mice and ctrl mice [42.9 (30.7-53.1) vs. 34.3% (31.2-53.3), p= 0.7; 52.4 (44.3-59.3) vs. 45.9% (41.2-60.2), p= 0.7, respectively] (Figure 3d). However, the proportions of CD3+CD8+ [31.5 (22.1-34.7) vs. 1.3% (0.6-5.4), p=0.001] and CD3+CD4+cells [39.7 (26.8-59) vs. 4.5% (2.6-10.5), p=0.007] were higher in EM mice. Conversely, CD3+CD4-CD8-cells were enriched in ctrl mice [82.3 (78.8-94.9) vs. 26.2% (12.9-35), p<0.0001] (Figure 3e) as well as CD3+CD4+CD25+ cells [24.9 (22-26.3) vs. 6.2% (4.1-6.8), p=0.0002] (Figure 3f), indicating the prevalence of T-lymphocytes with a pro-inflammatory phenotype in EM mice.

The transcript levels of several type I IFN-stimulated genes (i.e. IFIT3, IFI44, RSAD2) were two- to ten-fold higher (Figure 3g) in the gastrocnemius muscle of EM mice while transcript levels of several type II IFN-stimulated genes (PSMB8, GBP2, GBP1 and CXCL9) were increased by two- to seven-fold (Figure 3h).

Together, these results indicate that EM mice show a pro-inflammatory muscle environment that resembles IM regardless of disease subtype.

## Discussion

Previous attempts to overcome the technical limits of CIM using C-protein polypeptides have failed to reproduce the characteristics of IM in mice (19). Such polypeptides are located within the most immunogenic region according to Sugihara et al. (fragment 2, aa 284-580) (21).

Based on *in silico* immunogenicity prediction, we identified a polypeptide encompassing aa 965-991 of C-protein, which was synthetised with high purity. The polypeptide induced a reproducible experimental IM model (EM) by single immunisation in several independent cohorts of mice, recapitulating the hallmarks common to all IM subtypes at the clinical, histological, cellular and molecular levels.

EM has several advantages compared to CIM. Polypeptide production is low-cost and does not require specific laboratory facilities. This makes polypeptide production more affordable and reliable, and enables harmonisation of the results. Similarly to CIM, the C57BL/6N background facilitates the potential induction of EM in genetically-engineered mice which could prove valuable in fostering pre-clinical research in IM.

Using solely a polypeptide portion of the C-protein, we were able to confirm previous findings obtained in CIM mice including decreased exercise capacity as well as endomysial, perimysial and perivascular mononuclear cell infiltrates (T cells and macrophages) within the muscle (21). We additionally provide new insights in the characterisation of the model by demonstrating muscle weakness, increased serum CK levels, muscle fibre lesions (i.e., MHC-I expression, atrophy, mitochondrial abnormalities, sarcolemmal irregularities, sarcomeric disruptions and necrosis), immune cell compartment imbalance and IFN-signature (type 1 and 2) within the muscle. Muscle inflammation in CIM was previously shown to begin resolving after D28 while muscle function was not assessed at this timepoint (21). Of note, in the present study, clinical and biological phenotypes in EM persisted for at least 31 days, thus providing a larger experimental window.

Although the exact pathophysiological mechanisms of IM are unknown, data indicate that some mechanisms are shared by all IM subtypes (5), whereas others are subtype-specific (12). This observation probably explains that certain current therapeutic strategies improve all IM subtypes (i.e. corticosteroids, rituximab) while others are only effective in some subgroups (43). EM may represent an advantage in a strategy aimed at deciphering the underlying mechanisms and/or testing the efficacy of targeting common mechanisms of IM. However, this may also represent a limitation in a strategy aiming at deciphering subtype-specific aspects of the disease. For example, the specific role of auto-antigens and/or auto-antibodies is unlikely to be answered by EM, but possibly addressed by other animal models (e.g. based on immunization against specific auto-antigens (44) or on the transfer of specific auto-antibodies). In conclusion, EM mice are an optimized and characterised version of CIM that recapitulates common hallmarks of IM. This costless, high throughput, reliable and stable model, obtained in the most commonly used background for genetic engineering in mice (C57BL/6N), may foster a better understanding and pre-clinical treatment of IM.

## Acknowledgements.

We thank the IGBMC animal house facility as well as R. Lutzing, W. Magnant, A. Vincent and S. Falcone for their excellent technical assistance. We acknowledge the IGBMC Flow cytometry platform as well as C. Ebel and M. Philipps for technical support and helpful discussions. We thank the IGBMC peptides synthesis facility as well as P. Eberling for polypeptides synthesis.

We thank Mr Pierre Pothier for proofreading.

## Funding

This work was supported by French state funds from Agence Nationale de la Recherche ANR-GC-MYOS to A.M.

## Data availability

Data will be shared upon request from any qualified investigator.

## Conflict of interest

None declared.

---

**Supplementary Figure 1. Characteristics of three polypeptides from human MYPC fast type.** a) Peptide purity assessment by reversed-phase high-performance chromatography analysis. Coefficient of purity refers to the percentage of the target peptide compared to impurities that absorbs at 215 nm wavelength.

**Supplementary Figure 2. Characterisation of muscle impairment in mice treated with polypeptides of human MYPC fast type.** a) Representative images of H&E staining of gastrocnemius cryosections at D21 after immunisation with peptides 1 and 2 compared to mice treated only with phosphate-buffered saline (PBS, ctrl PBS) or with Freund’s adjuvant (ctrl FA). Scale bar: 50 µm. These experiments were performed in male mice. b) Footprint analysis in female mice immunised with peptide 3 at D20. The hind base width calculation is shown.

**Supplementary Figure 3. Muscle immune cells infiltration and myofibre abnormalities in female mice treated with polypeptides of human MYPC fast type.** a) Representative images of haematoxylin and eosin (H&E) staining of anterior tibialis, gastrocnemius and quadriceps cryosections at day 14 (D14), D21 and D31 after immunisation with peptide 3 in female mice. Scale bar: 50 µm.

**Supplementary Figure 4. Electron microscopy analysis in quadriceps muscle at D21.** Sarcolemmal invagination (red arrows) and mitochondrial fragmentation (orange arrows) are shown in peptide 3-treated mice compared to control mice (n= 3 for each condition). These experiments were performed in female mice.

**Supplementary Figure 5. Representative figure of the gating strategy performed at muscle flow cytometry in experimental myositis mice.**

**Supplementary Table 1.**
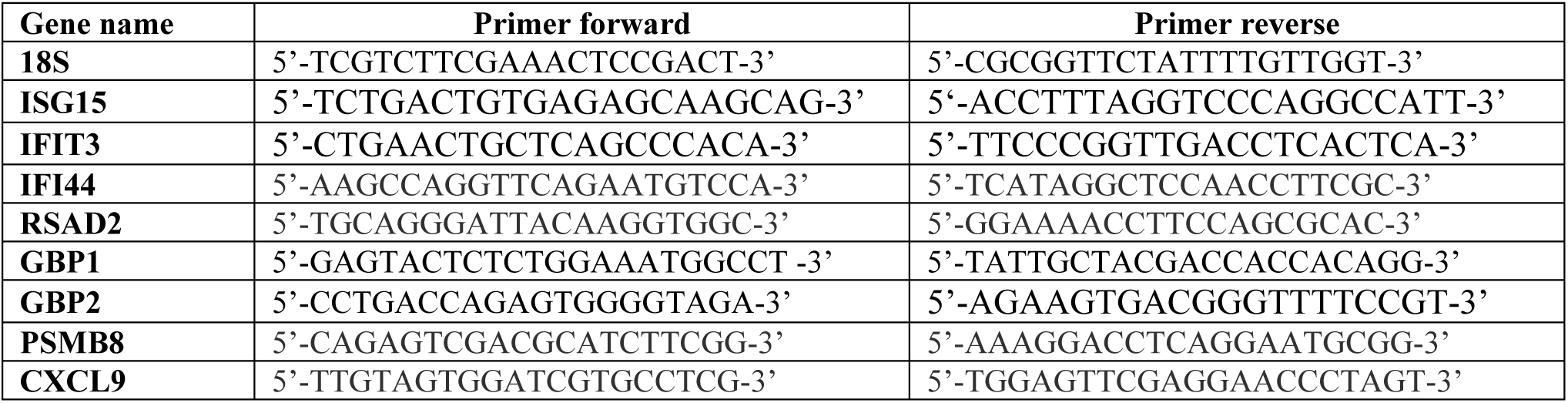
Sequence of gene-specific primers used for quantitative polymerase chain reaction.

**Supplementary Table 2.** List of autoantibodies used for muscle flow cytometry experiments.

| ANTIBODIES | SOURCE | IDENTIFIER | DILUTION |
| --- | --- | --- | --- |
| CD45 Alexa Fluor 700 (clone: 30-F11) | BioLegend | 103128 | 1:200 |
| CD3 PerCPCy5,5 (clone: 145-2C11) | BioLegend | 100328 | 1:25 |
| CD4 PE (clone: RM4-5) | BD Pharmigen | 553049 | 1:200 |
| CD8 Brilliant Violet 785 (clone: 53-6.7) | BioLegend | 100749 | 1:200 |
| CD25 Brilliant Violet 650 (clone: PC61) | BioLegend | 102038 | 1:100 |
| CD11b Pe-Cyanine7 (clone: M1/70) | eBioscience | 25-0112-82 | 1:200 |
| F4/80 APC-e-Fluor780 (clone: BM8) | eBioscience | 47-4801-82 | 1:200 |
| Ly6C PE-CF594 (clone: AL-21) | BD Biosciences | 562728 | 1:400 |
| CD206 FITC (clone: C068C2) | BioLegend | 141704 | 1:100 |
| Annexin V FITC | BD Biosciences | 560931 | 1:100 |

## Notes

### Competing Interest Statement

The authors have declared no competing interest.

