## Supplementary figures and images for "Experimental myositis: an optimised version of C-protein-induced myositis"

### Supplementary Figure 1

a

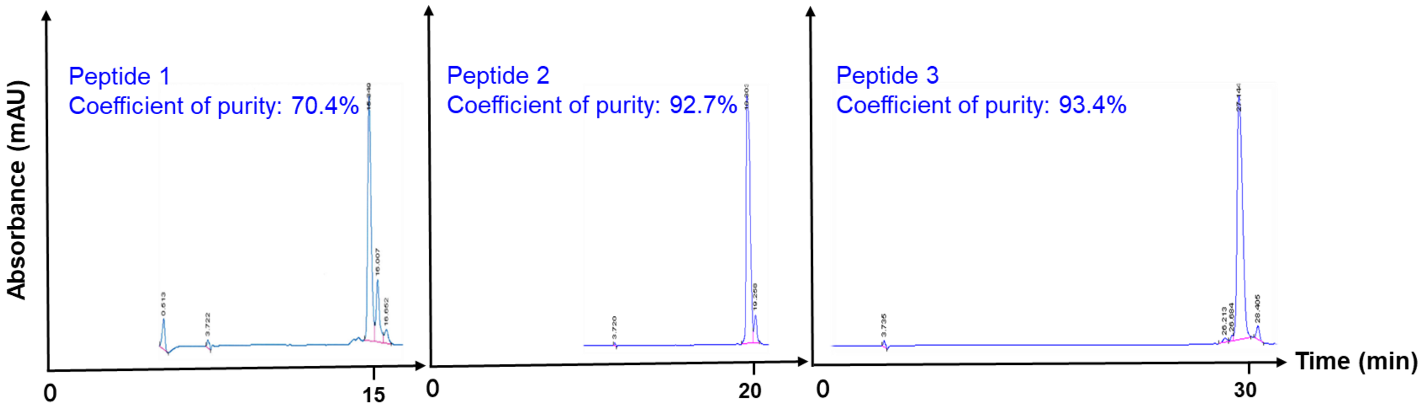


supplementary figure 1

### Supplementary Figure 2

**a**

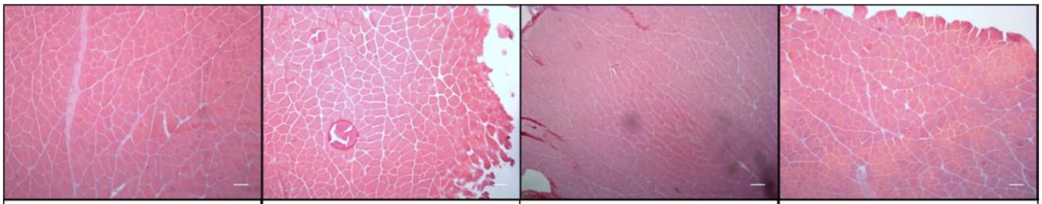

| ctrl PBS | ctrl FA | peptide 1 | peptide 2 |
|---|---|---|---|

**b**

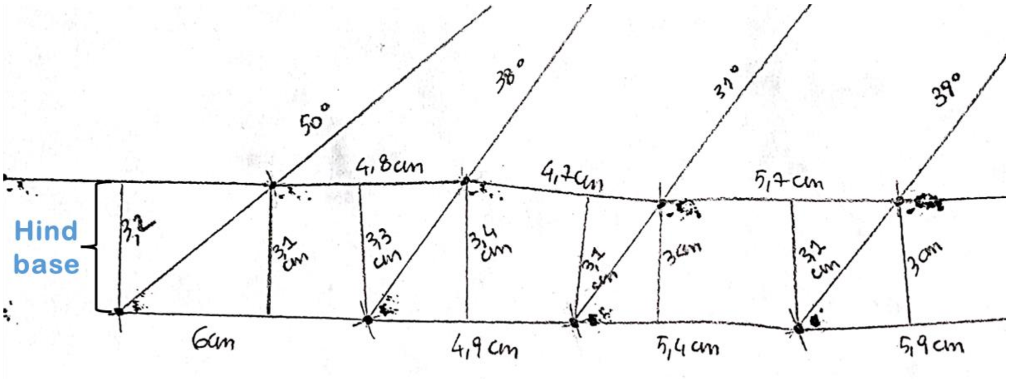


**supplementary figure 2**

### Supplementary Figure 3

**a**

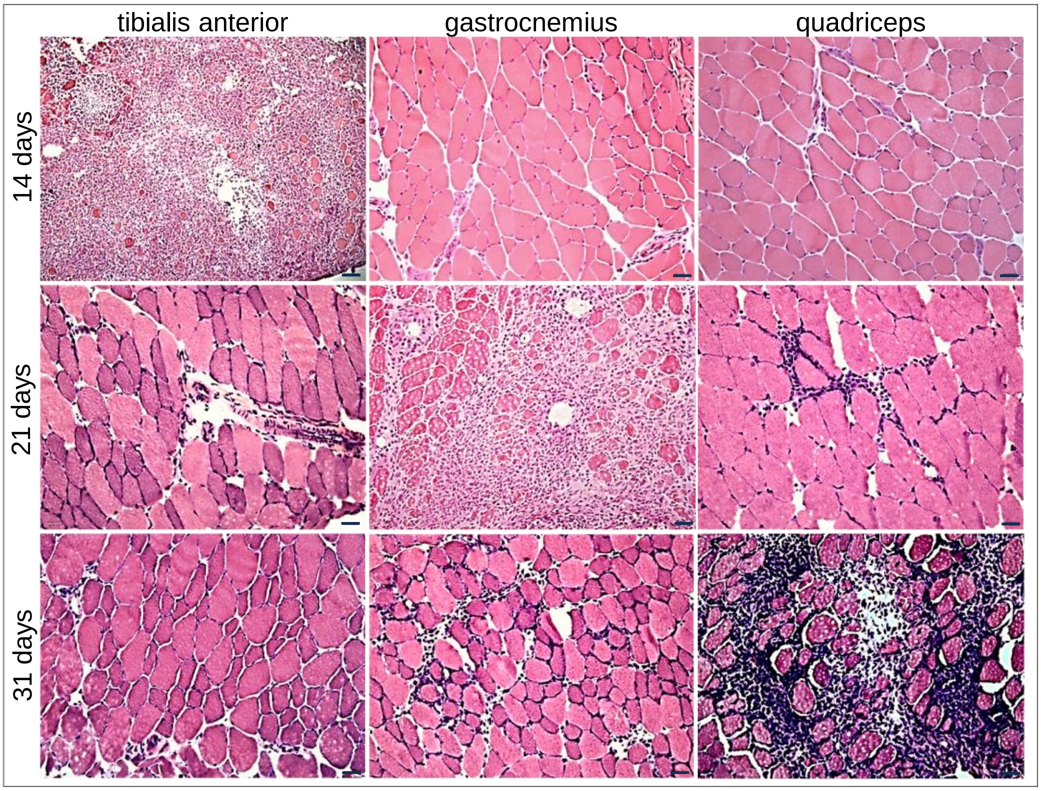


**supplementary figure 3**

### Supplementary Figure 4

a

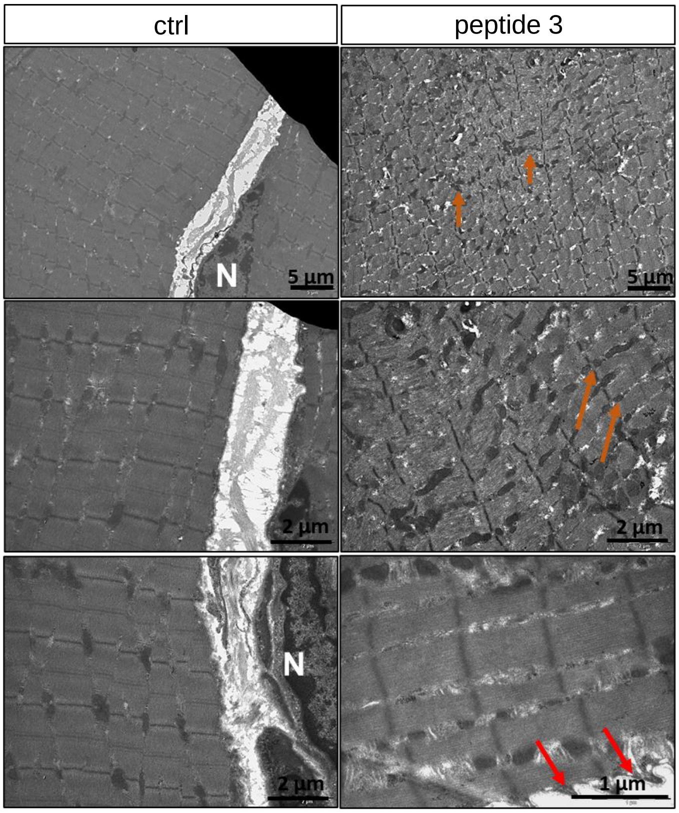


**supplementary figure 4**

### Supplementary Figure 5

**a**

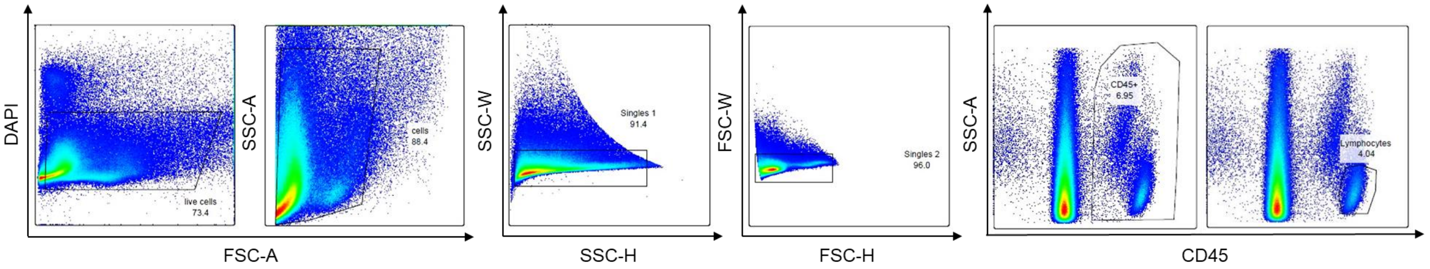


**supplementary figure 5**
